# Systems biology analysis of publicly available transcriptomic data reveals a critical link between *AKR1B10* gene expression, smoking and occurrence of lung cancer

**DOI:** 10.1101/759282

**Authors:** Juan M. Cubillos-Angulo, Eduardo R. Fukutani, Luís A. B. Cruz, María B. Arriaga, João Victor Lima, Bruno B. Andrade, Artur T. L. Queiroz, Kiyoshi F. Fukutani

**Author notes:** **Corresponding Authors** (BBA), (ATLQ) and (KFF). BBA, ATLQ and KFF equally contributed to the work.

## Abstract

**Background:** Cigarette smoking is associated with increased risk of developing respiratory diseases and various types of cancer. Early identification of such unfavorable outcomes in patients who smoke is critical for optimizing personalized medical care.

**Methods:** Here, we perform a comprehensive analysis using Systems Biology tools of publicly available data from a total of six transcriptomic studies, which examined different specimens of lung tissue and/or cells of smokers and nonsmokers to identify potential markers associated with lung cancer.

**Results:** Expression level of twenty-two genes was capable of classifying smokers from non-smokers. A machine learning algorithm revealed that *AKR1B10* was the most informative gene among the 22 DEGs accounting for the classification of the clinical groups. *AKR1B10* expression was higher in smokers compared to non-smokers in datasets examining small and large airway epithelia, but not in the data from a study of sorted alveolar macrophages. We next tested whether *AKR1B10* expression could be useful in identification of cancer tissue in patients who were not exposed to smoking. *AKR1B10* expression was substantially higher in lung cancer specimens compared to matched healthy tissue obtained from nonsmoking individuals (accuracy: 80%, p<0.0001). Finally, we searched the expression of 11 single nucleotide polymorphisms (SNPs) of *AKR1B10* worldwide. We found that the SNP rs782881 was the most frequent mutant allele in the majority of continents. Africa was the continent which exhibited higher frequency of SNPs associated with lower *AKR1B10* expression and displayed lower lung cancer incidence and deaths attributable to tobacco.

**Conclusion:** The systematic analysis of transcriptomic studies performed here revealed a potential critical link between *AKR1B10* expression, smoking and occurrence of lung cancer.

## Introduction

Worldwide, cigarette smoking is a life-style habit of approximately 1.1 billion individuals and is associated with more than six million deaths annually [1]. The immunological responses in persons chronically exposed to smoke from cigarette are characterized by protracted secretion of inflammatory factors and by accumulation of several leukocytes in lung tissue and production of pro-fibrotic mediators such as transforming growth factor (TGF)-β [2, 3]. These inflammatory perturbances likely result in increased risk development of tobacco associated morbidity including several types of cancer [4], autoimmune disorders [5], chronic obstructive pulmonary diseases[6] and respiratory infections [7].

The role of tobacco smoking in the induction of disturbances in cell/tissue homeostasis and gene mutations, broadly or specifically associated with several types of tumors, have been investigated. Smoking-related malignancies have been reported to be associated with DNA methylation [8] and mutations in several proto-oncogenes, such as *p53, KRAS, BRCA-1, BRCA-2, GPX2, GABP, TCF3, CRX, CYP2A13, CYP2A6, CYP2B6,* among others [9–12]. In addition, it has also been reported that components of cigarette smoking modulate immune cell functions, which could lead to loss of T-cell proliferation and antibody responses [13]. Furthermore, chromosomal instability, epigenomic alterations and several mutations have been reportedly associated with lung cancer in particular [14]. Thus, in general, all of these events ultimately culminate with altered gene expression, even though the conversion of carcinogens to DNA adducts is more efficient in some individuals than in others [15]. Therefore, understanding the expression of these gene is important to fully understand the link between smoking exposure and risk of cancer development.

Identification of genetic markers predictive of cancer development is of utmost importance for promoting personalized medicine [16]. Such markers could be implemented as screening strategy for patients who exhibit strong risk factors for cancer, such as cigarette smoking. To identify such potential markers, we performed a systematic analysis of publicly available data from transcriptomic studies performed in lung tissue and/or cells and found that, among most of the studies investigated, increased expression of the gene *AKR1B10* was strongly associated with cigarette smoking as well as lung cancer. Development of a point-of-care assay to assess *AKR1B10* expression in individuals exposed to cigarette smoking may serve as a relevant tool to identify those with high risk of cancer.

## Methods

### Ethics Statement

There were no patients directly involved in the research. The present study used publicly available gene expression data from previous studies to perform a meta-transcriptome analysis. All information given to the research team were de-identified.

### Description of discovery datasets

We accessed the Gene Expression Omnibus (GEO-NCBI – https://www.ncbi.nlm.nih.gov/geo/) and have looked for datasets with tobacco smoking information in human biopsies or tissues, without diagnosis of other comorbidities. Five datasets have been found in GEO. Thus, two previously published microarray datasets were selected to be used as a discovery set (available from the GEO under accession no. GSE4498 [17] and GSE3320 [18]) and 3 have been used as validation set (GSE20257 [19], GSE17905 [20] and GSE13931 [21]). The Dataset GSE4498 [17] was designed with samples of human small airway bronchial epithelium of smokers (n=10) compared to matched samples from non-smokers (n=12). The dataset GSE3320 [18] was extracted from samples of human small airway bronchial epithelium to assess gene expression in phenotypically smokers (n=6) compared to matched non-smokers (n=5). These included datasets using the same method to collect the samples, by fiberoptic bronchoscopy and brushing. In addition, these studies used a similar transcriptional protocol using the platform Affymetrix Array, making possible to combine both datasets in a discovery set.

### *In silico* validation

We next performed validation of the differentially expressed genes detected in the first phase of the investigation using three distinct datasets selected by examination of gene expression by smoking status: (i) GSE20257 was published by Shaykhievet a l[19]. In this study, they used samples of small airway epithelium collected from individuals who were smokers (n=51) and also from those who did not smoke (n=42) and performed an analysis of microarray assays in theses samples. (ii) GSE17905 was published by Wang et al [20]. The authors used large airways samples collected by bronchoscopy of 31 smokers and 21 non-smoker individuals and also performed a microarray analysis. (iii) GSE13931 was published by Carolan et al [21]. The investigators used alveolar macrophages collected by broncho-alveolar lavage of 30 smokers and 19 non-smokers and performed a microarray analysis. (iv) Finally, GSE19804 was available in a publication from Lu et al [22]. This dataset had information of gene expression (assessed by microarray) of sixty (n=60) pairs of lung cancer tissue and adjacent normal lung tissue from female patients who were not exposed to cigarette smoking.

The datasets were obtained using the *GEOquerry* [23] package and raw expression data of 22 samples present on GSE4498 and 11 samples on GSE3320 were normalized and log2 transformed by *preprocesscoreR* package [24]. Duplicated probes were collapsed by *collapserows* function in *WGCNA* package [25] and all common genes to both datasets were kept and used to merge the datasets. The expression data was submitted to a correction procedure of batch effect using an empirical Bayes framework implemented in the *COMBAT* function available in *SVApackage* [26].

### Statistical analysis

The DEGs were identified by applying the absolute ≥1.0 log2-fold-change threshold and p-value corrected with FDR adjustment for multiple testing (FDR = 5%), from *limmapackage* [27]. A volcano plot we used to identify changes in gene expression, the significance versus fold-change on the y and x axes, respectively. We use Venn diagrams to visualize all possible logical relations between all the DEGs between smokers and nonsmokers in all datasets evaluated. The modular analysis was performed using the *Cemitool* package [28]. It is based on Weighted correlation network analysis (WGCNA) and default parameters was employed (Beta Parameter = 7). The module annotation was performed with the Kegg database v6.2 [29] and Gene Set Enrichment Analysis (GSEA) algorithm is available internally in the *Cemitool* package and the Single sample Gene Set Enrichment analysis (ssGSEA) was performed with *GSVA* package [30]. The significant and annotated pathways were clustered using Euclidean distance as dissimilarity measure and average linkage for between-cluster separation *(hclust* function in the stats package in R 3.2.2). All The heatmap was generated in R via the *heatmap.2* function from the *gplots* package, using the “scale = “row” switch to Z-score standardize the rows [31]. PCA was performed in order to compare and visualize the expression values of all genes to estimate the variance of the global gene expression with the function *prcomp* a native package in R. The decision trees were employed to validate and identify the minimal gene set that correctly classifies the smokers from nonsmokers from the 22-gene signature [32]. To estimate the decision tree models accuracy, we performed a 10-fold cross validation. The partition procedure was applied to avoid bias in the training/test sets sampling. Thus, the training set was used to tune the parameters, learning and building a model. The validation set was used to test the classifier performance. The sensibility and specificity were measured from the confusion matrix and visualized in the receiver–operating characteristic curve (ROC) [32]. Accuracy was evaluated by area under the curve of ROC plot. We used the genotype-tissue expression project (GTEx) test association of the SNPs in the *AKR1B10* gene with tissues of the respiratory tract [33].

## Results

### Meta-transcriptome signature of smoking

After preprocessing and merging the datasets, we applied a Principal component analysis (PCA) algorithm using the expression values of all genes to estimate the variance of the global gene expression. This analysis revealed that the subgroups of smokers and non-smokers could not be separated, and two main groups containing both smoker and non-smoker individuals were observed (Figure 1A). To visualize the overall profile of individual gene expression, we used a volcano plot (Figure 1B). This approach indicated presence of a total of 800 statistically significant genes (p<0.05, corrected by Benjamini–Hochberg false discovery rate [FDR]), of which 375 genes were upregulated and 425 genes were downregulated (Figure 1B). Additional analyses identified 22 the differentially expressed genes (DEGs), defined here and genes which exhibited more than ± 1 fold-difference variation (smokers vs. non-smokers) and a significant p-value after FDR adjustment (p<0.05). Such DEGs were inputted in an unsupervised two-way hierarchical clustering analysis. The results demonstrated that when considered together, the 22-gene signature was capable of classifying smokers from non-smokers into completely separate clusters (Figure 1C).

**Figure 1.**
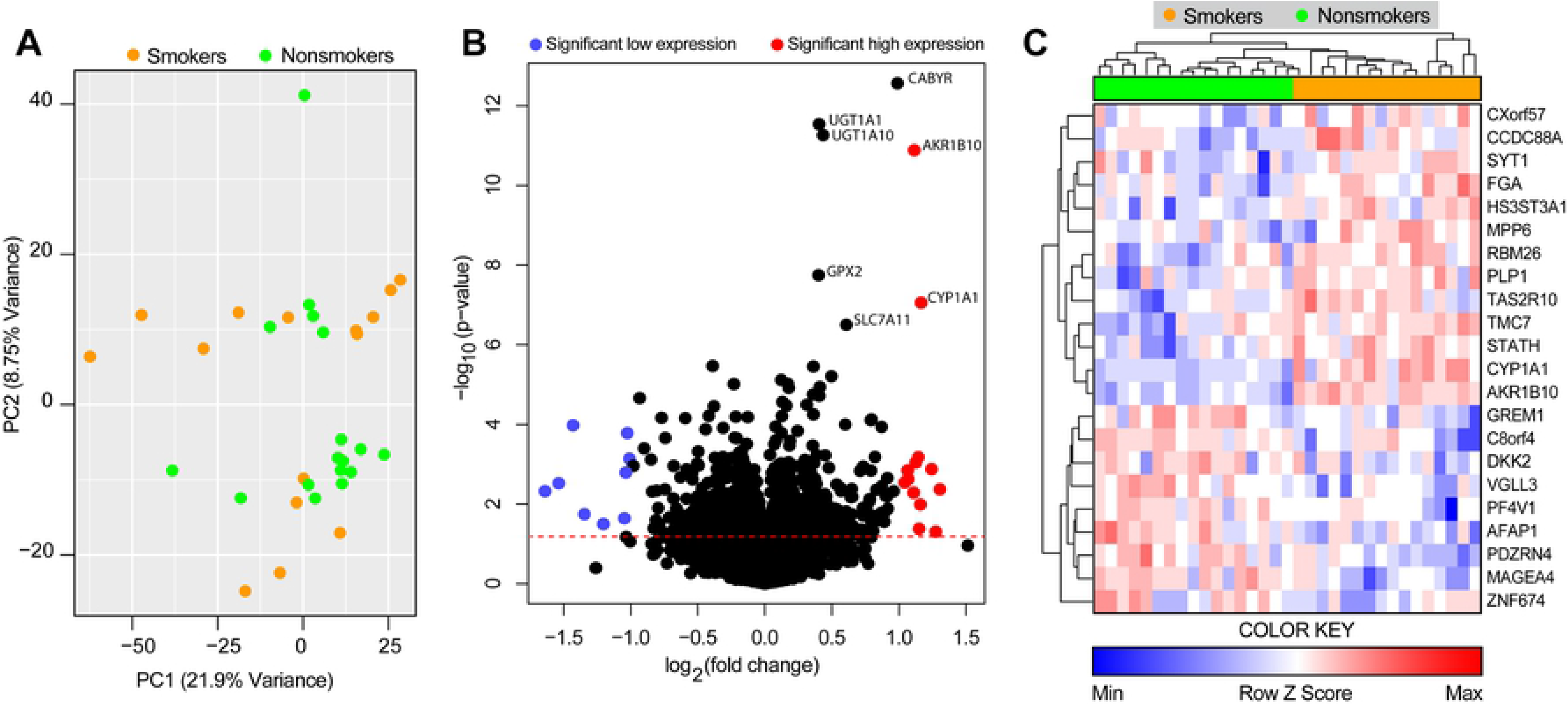
Differentially expressed genes associated with cigarette smoking. We analyzed publicly available data of two datasets of small airways transcriptome (RNAseq). **(A)** A principal component analysis (PCA) model of 13,516 genes was used to distinguish smokers from nonsmokers. **(B)** Volcano plot of all genes (smoker vs. nonsmokers). **(C)** 22 differentially expressed genes (DEGs), defined as p<0.05 after 1%FDR and 1.0-fold change expression, were found and together were able to discriminate the clinical conditions.

To delineate the gene pathways from which the overall transcription profile in smokers vs. non-smokers were involved, we used the *CemiTool* package[28]. We detected three distinct co-expressed gene modules, annotated in Kyoto Encyclopedia of Genes and Genomes *(Kegg)* database (Module [M] 2, M7, M14). Two modules were enriched in the non-smoking samples compared to smokers, based on the normalized enrichment scores (NES) (Figure 2A). The first module (M2), found to be enriched in non-smokers was Glycosaminoglycan biosynthesis chondroitin (log10 p = 1.57). A second module (M7) was overrepresented in smokers compared to non-smokers and showed to be enriched in the Peroxisome proliferator-activated receptor (PPAR) signaling pathway (log10 p-value = 1.5). A third module, also more representative in smokers encompassed colorectal cancer (p< 5.34e-38) and basal cell carcinoma (log10 p-value =3.2). We next calculated the NES for each top ranked pathway identified per individual study subject and found that, when considered together, such pathways were not able to cluster smokers and non-smokers separately (Figure 2C).

**Figure 2.**
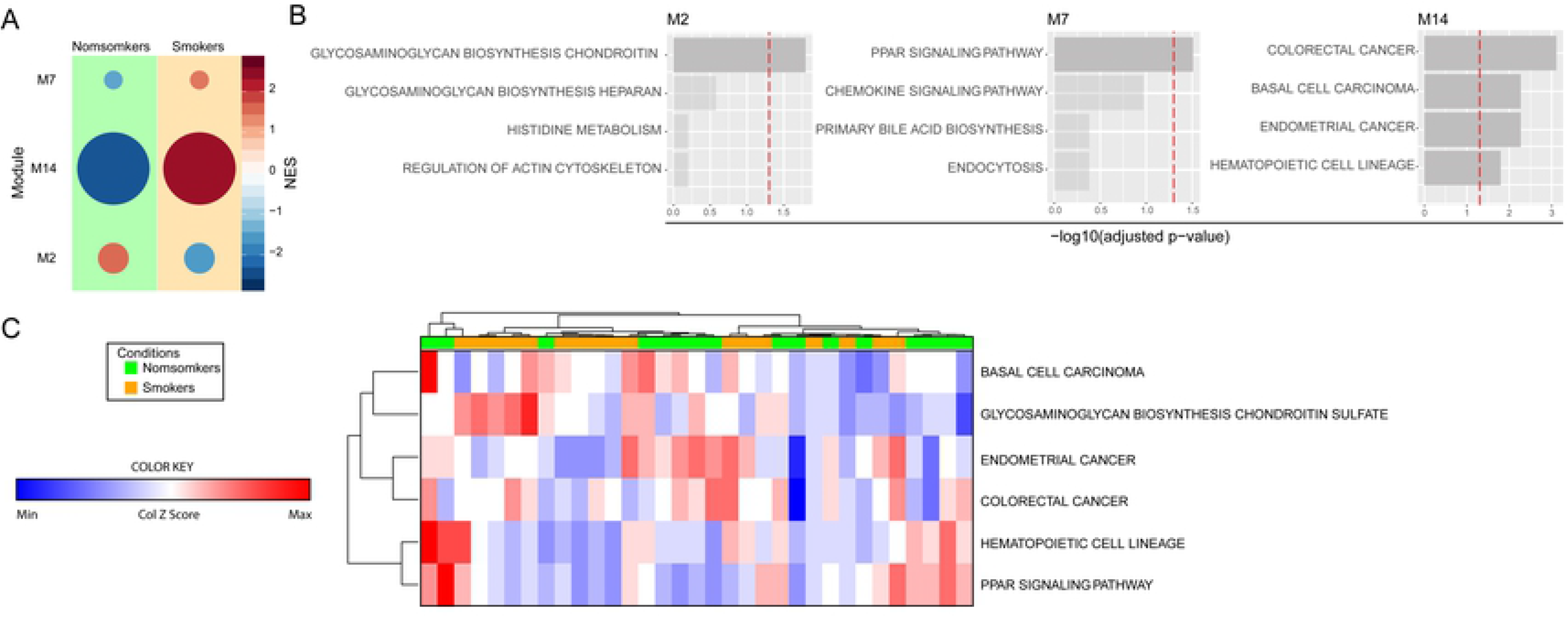
Gene pathway analysis in smokers and nonsmokers. **(A)** Co-expressed modules of all genes. Circle sizes are proportional to the normalized enrichment scores (NES). **(B)** The modules were annotated using Keg package for R. Dashed lines represent significance threshold. **(C)** Hierarchical cluster analysis (Ward’s method) using the NES scores for each annotated module and calculated for each person was employed test discrimination between smokers and nonsmokers.

### *AKR1B10* gene expression level in lung tissue, but not in alveolar macrophages, exhibits high accuracy in discriminating smoking from nonsmoking individuals

To validate our discoveries, we tested the 22 DEGs identified in our analyses in 3 distinct datasets that compared smokers and non-smokers: (i) GSE20257, that was composed by data from small airways samples, (ii) GSE17905, which compared gene expression from large airways samples and (iii) GSE13931, which used data from alveolar macrophages. Discriminant analyses using Receiver Operating Characteristic (ROC) curves were able to reveal high accuracy of such gene signature to distinguish smokers from nonsmokers in the two datasets that large and small airway samples (GSE20257 Area under the curve [AUC]: 0.862, p<0.0001; GSE17905 AUC: 0.864, p<0.0001). The same approach indicated that when a dataset from alveolar macrophages was considered, the 22-gene signature was not able to distinguish the study groups (GSE13931AUC: 0.607, p=0.150) (Figure 3A). We next employed a machine-learning approach using decision trees to identify which markers from the 22-gene signature would exhibit more robust discrimination power in each dataset evaluated. Of note, the gene *AKR1B10* was the most informative gene in the discovery set and also in the two distinct datasets that used large or small airway tissue (Figure 3B). In the dataset that used gene expression values form alveolar macrophages, *AKR1B10* was not shown to be relevant in discrimination, and a combination of two other genes *(VGLL3* and *TAS2R10*) accounted for the differences between smokers and non-smokers (Figure 3B). *AKR1B10* expression was higher in smokers compared to non-smokers in all datasets evaluated, except again in the GSE13931, which used data on alveolar macrophages (Figure 3C). Furthermore, we plotted Venn diagrams of all the DEGs between smokers and non-smokers in each dataset to verify overlaps. We confirmed that *AKR1B10* was a DEG commonly shown in the discovery set as well as in the databanks which used airway tissue samples, but not in the alveolar macrophage dataset (Figure 3D). The two other DEGs found in smokers were *CYP1A1* and *HS3ST3A1* (Figure 3D). *CYP1A1* encodes a protein that localizes at the endoplasmic reticulum and its expression is induced by polycyclic aromatic hydrocarbons, some of which are found in cigarette smoke [34]. *HS3ST3A1is* a member of the heparan sulfate biosynthetic enzyme family [35].

**Figure 3.**
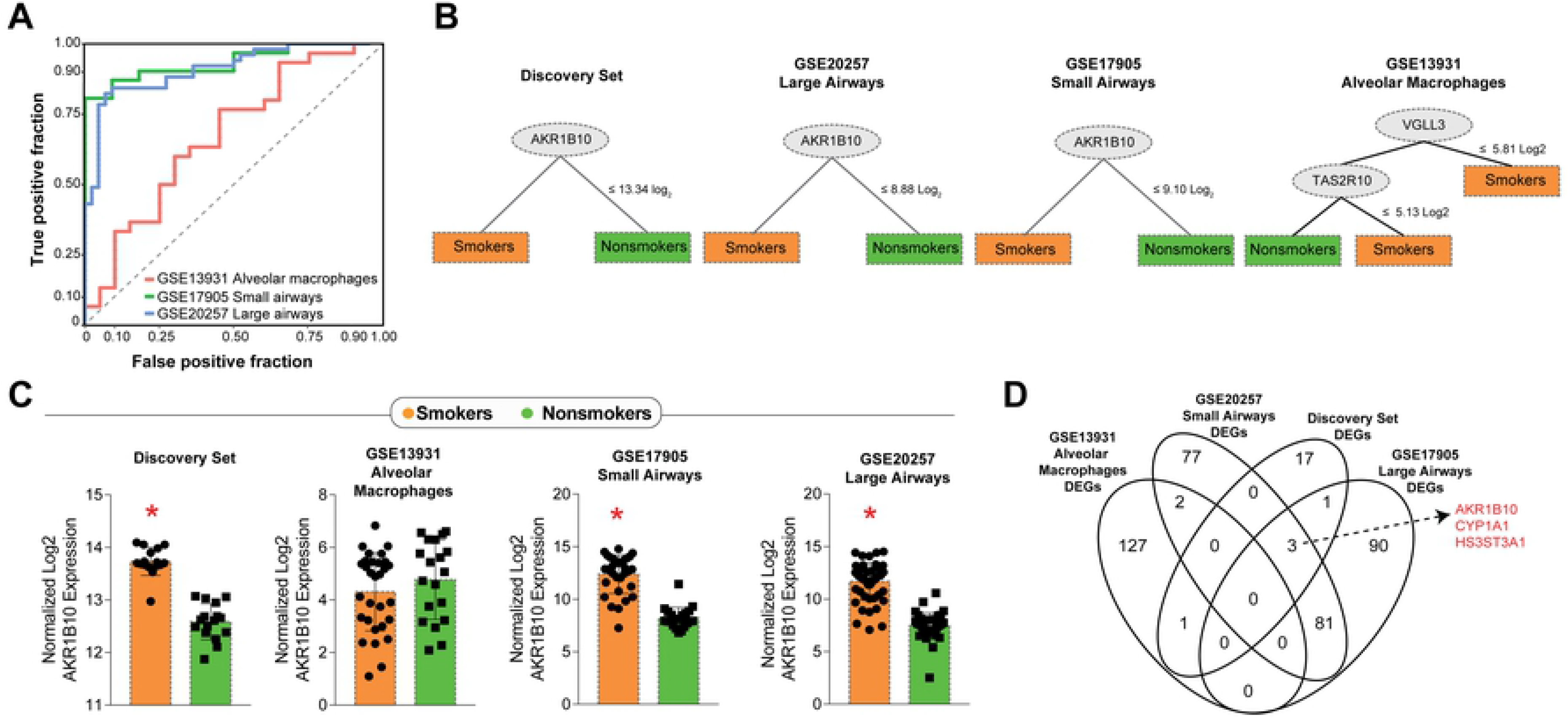
Defining the molecular signatures of smoking. **(A)** Data on the 22 DEGs found in our discovery analyses were used to validate discrimination between smokers and nonsmokers in 3 different previously published datasets. **(B)** Machine-learning decision trees were built for each dataset to describe the most relevant genes driving discrimination. Of note, the gene AKR1B10 was found to be the main discriminator in 3 out of the 4 datasets examined. **(C)** Scatter plots of the AKR1B10 gene expression in the 4 datasets. **(D)** Venn diagram of the DEGs in each dataset shows AKR1B10 in the intersection of 3 datasets extracted from lung tissue specimens but not included among DEGs from alveolar macrophages. *p<0.05 (Student’s t-test).

### *AKR1B10* is able to discriminate lung cancer patients who do not smoke

The results described above demonstrate that higher *AKR1B10* expression hallmarks tissue airways from smokers. Given that smoking is a well-established risk factor of lung cancer[9], we hypothesized that *AKR1B10* gene expression could also be useful in identification of cancer patients who were not exposed to smoking. To test this hypothesis, we downloaded the dataset GSE19804, which included tissue samples from non-small cell lung cancer as well as ipsilateral healthy lung tissue obtained from patients who did not present history of cigarette smoking. The *AKR1B10* gene expression was substantially higher in the specimen collected from the tumor compared to the healthy lung tissue in the same patients (Figure 4A). ROC curve analysis indicated that *AKR1B10* gene expression value was able to discriminate cancer and non-cancer tissue with relative high accuracy (AUC 0.80, p<0.0001) (Figure 4B).

**Figure 4.**
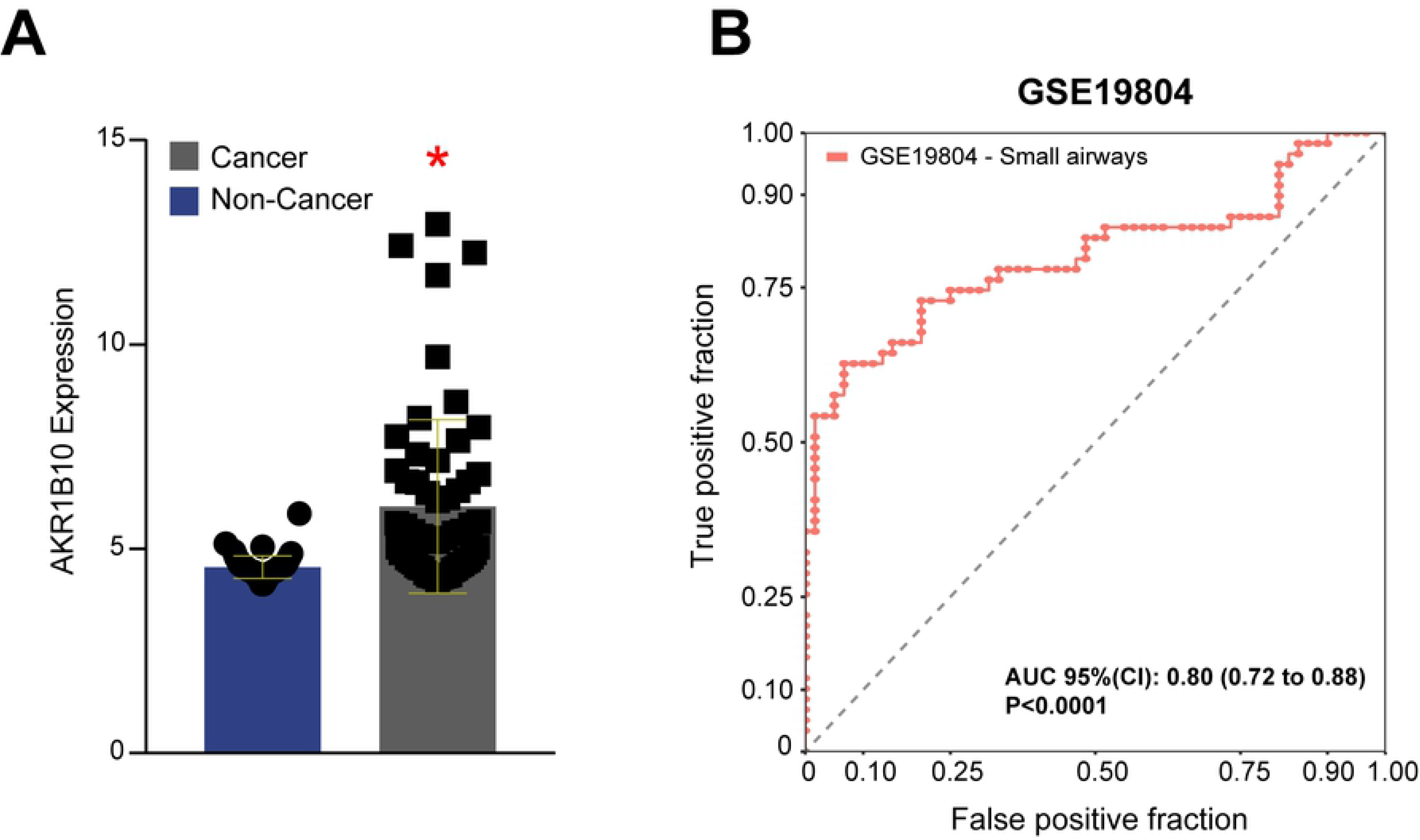
In nonsmokers, higher AKR1B10 expression is a biomarker of lung cancer. **(A)** We analyzed AKR1B10 gene expression values in a published dataset of neoplastic lung tissue microarray in nonsmoking individuals who were diagnosed with lung cancer and compared to ipsilateral healthy lung tissue specimens (controls.) Scatter plots of AKR1B10 gene expression in the groups. *p<0.05 (Student’s t-test). **(B)** Receiver Operator Characteristics (ROC) indicated a high accuracy to discriminate cancer tissue from controls.

### Distribution of Single Nucleotide Polymorphisms of AKR1B10, of lung cancer incidence and of deaths attributed to tobacco worldwide

We next searched for all the single nucleotide polymorphisms (SNPs) of *AKR1B10* that have been published so far and found 11, which have been specific in the gene region (Figure 5A). The allele I was denominated the normal nucleotide whereas allele II was the SNP. In Europe, Asia and in the American continents, the most frequent mutant allele was rs782881 (48%, 54% and 42% of the population from these continents, respectively) (Figure 5A). In Africa, the most common mutant allele was rs782510 (78% of the population Figure 5A). Of note, analysis of two SNPs rs782510 and rs782881 with *RegulomeDB* [36]revealed that both SNPs are likely to affect binding and linked to expression of a gene target e-QTLs (expression quantitative trait loci).

**Figure 5.**
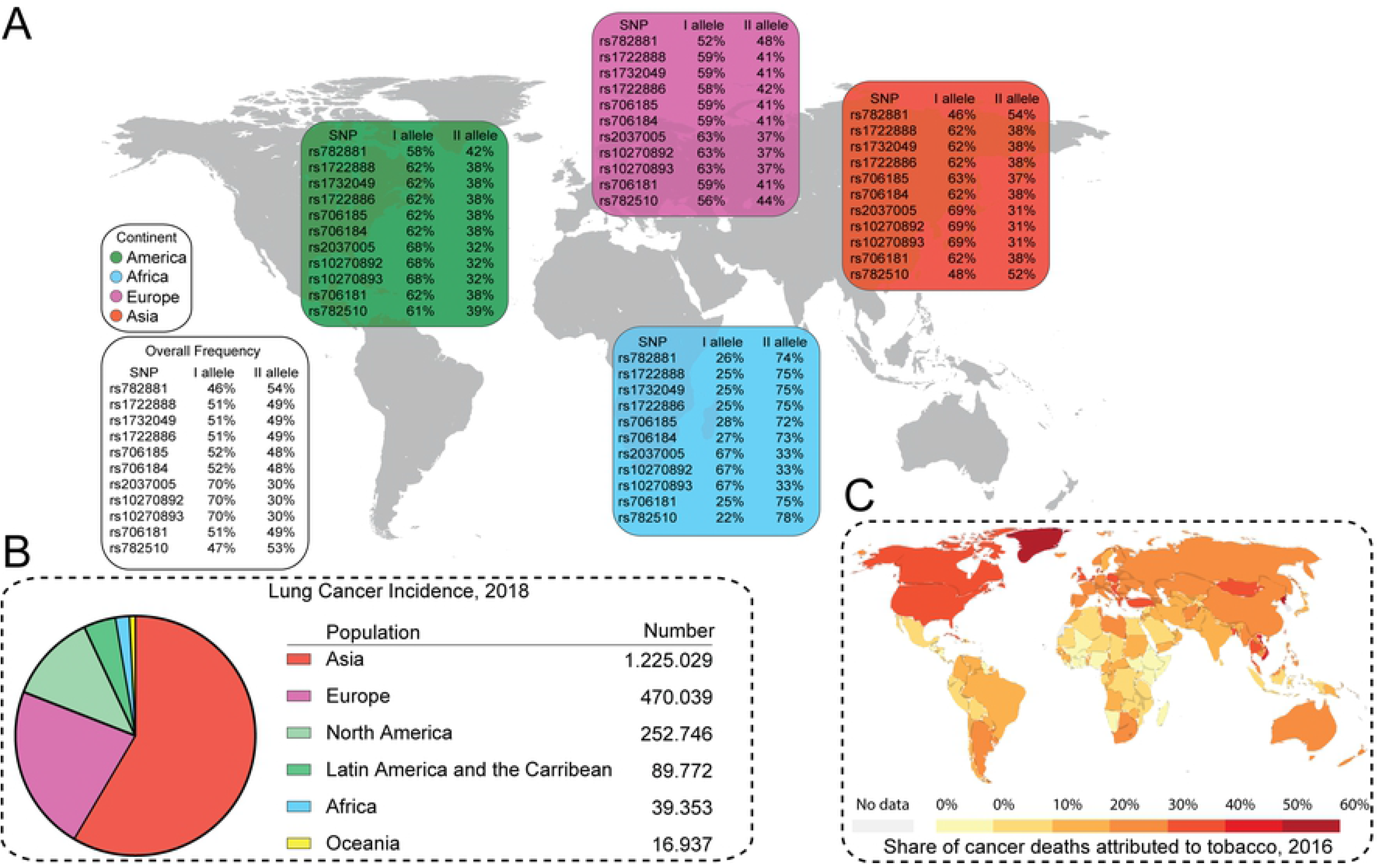
Allele frequencies of Single Nucleotide Polymorphisms of AKR1B10 and distribution of lung cancer incidence and of deaths attributed to tobacco worldwide. **(A)** Single nucleotide polymorphisms (SNPs) of the AKR1B10 which were published in the Genotype-Tissue Expression (GTEx) project. The allele I was denominated the normal nucleotide whereas allele II was the SNP. **(B)** Lung cancer incidence in both sexes obtained from the Global Cancer Observatory in 2018. **(C)** Graph obtained from “Our World in Data” which depicts cancer deaths attributable tobacco in 2016.

Importantly, when lung cancer incidence in 2018 in both sexes was ranked by continent, we observed that Asia accounted for the majority of cases, followed by Europe and North America, whereas incidence of lung cancer in the African continent was relatively low (Figure 5B). Finally, while assessing the Global Burden of Disease dataset [37]we found that these three continents had increased share of cancer deaths attributable tobacco whereas this variable was less representative in the African continent (Figure 5C).

## Discussion

In the present study, we examined a number of publicly available transcriptome data to identify a gene signature that could distinguish lung tissue specimens from smokers vs. non-smoking individuals. The most relevant finding of the initial part of analyses was the *AKR1B10* expression level was the most informative in such discrimination among three different datasets obtained from lung tissue, but not in the transcriptome data originated from alveolar macrophages. Moreover, *AKR1B10* gene expression level was able to reliably identify neoplastic from healthy lung tissue in persons not exposed to cigarette smoking. These findings are important because they have identified *AKR1B10* as a reliable biomarker which expression is triggered by cigarette smoking and is simultaneously observed in lung cancer specimens. It is possible that such gene may be involved in carcinogenesis associated with cigarette smoking. In fact, among the multiple carcinogens from cigarette smoke, the nitrosamine 4-(methylnitrosamino)-1-(3-pyridyl)-1-butanone (NNK) is described to play a critical role in lung carcinogenesis [38]. Carbonyl reduction takes place in both microsomal and cytosolic fractions from different human tissues such as lung and liver [39]. Within these subcellular fractions, several enzymes have been described to mediate NNK reduction, including the protein encoded by *AKR1B10,* which is from the aldo-keto reductase superfamily (AKR) [40]. Our findings not only confirm an association between *AKR1B10* and occurrence of lung cancer but also argues that expression of such gene may occur before onset of neoplastic lesions in smokers and may be suitable to be used for assessment of risk of cancer in highly exposed individuals. Additional studies are warranted to directly test this hypothesis.

The gene *AKR1B10* found differentially expressed in lung tissue from smokers vs. non-smokers has been previously described in experimental studies to play an important role in the pathophysiology of lung cancer [41]. *AKR1B10* is a regulator of the synthesis of fatty acid and participates in the metabolic pathway of lipids and isoprenoids [42]. In addition, the protein encoded by *AKR1B10* exhibits a high retinaldehyde reductase activity [43]. Importantly, *AKR1B10* can metabolize specific substrates, such as aldo-ketoreductases; farnesal, geranylgeranil, retinal and carbonyls[44]. Such activity is associated with promotion of carcinogenesis [45]. *AKR1B10* has also been shown to promote cancer cell survival by two distinct studies [46, 47]. These previous investigations revealed that knocking down *AKR1B10* expression induces cancer cell apoptosis and inhibited cancer cell proliferation, suggesting *AKR1B10* could serve as a potential therapeutic target.

Aside from being associated with lung carcinogenesis, *AKR1B10* expression has also been linked to development of several additional types of cancers. In hepatocellular carcinoma (HCC), *AKR1B10* expression is found upregulated, and experimental deletion of such gene inhibited the proliferation of HCC cells tumor growth in a xenograft mice model [48]. In HCT-8, a human colon adenocarcinoma cell line, and NCI-H460, a human lung carcinoma cell line, *AKR1B10* gene deletion has been shown to induce cell apoptosis and mitochondrial degeneration, leading to oxidative stress [46]. Furthermore, higher *AKR1B10* expression has been observed in squamous cell lung carcinoma (SCC) associated with smoking [49]. Finally, our findings indicate that *AKR1B10* is overexpressed in lungs of healthy people who smoke but had no cancer as well as in lung carcinoma from non-smokers. These observations argue that cigarette smoking already modifies the microenvironment of the lung epithelium probably creating a favorable scenario for carcinogenesis. This idea corroborates with previously published studies which demonstrated that smoking per se mediates upregulation of *AKR1B10* expression in the airway epithelia of healthy smokers with no evidence of lung cancer [50]. Thus, there is strong evidence to suggest that cigarette smoking-induced upregulation of *AKR1B10* may represent an initial critical step in the cascade of events leading to lung cancer.

In addition to *AKR1B10,* our analyzes revealed that two additional genes, *CYP1A1* and *HS3ST3A1,* overlapped in the datasets as DEGs capable of discriminating smokers from nonsmokers. Of note, *CYP1A1* has also been described to induce carcinogenesis, by promoting CYP-catalyzed epoxidation reactions, resulting in the formation of reactive metabolites that can cause DNA [51, 52]. Moreover, *CYP1A1* polymorphisms in smokers increase susceptibility to stomach cancer [53]. Furthermore, *HS3ST3A1* gene encodes the enzyme 3-O-sulfotransferase, which catalyzes the biosynthesis of a specific subtype of heparan sulfate (HS), 3-O-sulfated heparan sulfate, which is found to be upregulated in human lung cancer specimens and to contribute to its elevated metastatic potential [35]. Thus, the three genes found commonly differentiate regulated in individuals exposed to cigarette smoking are all know to favor development of cancer and could be used as early biomarker of disease progression in high risk populations, but future studies specifically designed to test this hypothesis are necessary.

Our investigations ultimately demonstrated that the AKR1B10SNP rs782881, which is linked to potential increase in gene expression, is the most frequent in Europe, Asia and American continents, where the incidence of lung cancer is high and cigarette smoking is also more frequent. The presence of this SNP has been correlated with increased number of cigarettes consumed per day [54]. Noteworthy, the African continent has much lower frequency of SNP rs782881 and coincidently a relatively lower incidence of lung cancer and consumption of cigarettes compared to the other continents. These findings make us hypothesize that there are SNPs in the *AKR1B10* gene that may protect against lung cancer. It would be important to conduct studies directly testing the association between different *AKR1B10* SNPs and occurrence of lung (or other) cancers in distinct populations worldwide.

Our study has several strengths such as the large number of samples evaluated, the use of discovery and validation datasets using different lung tissue/cellular types and different clinical conditions, as well as the detailed search of information about the distribution of different *AKR1B10* SNPs in the world. An important limitation was the low number of studies included, which was dependent of publicly available datasets. In addition, we have not performed validation in experimental systems. Regardless, by performing a systematic analysis of publicly available data from transcriptomic studies of lung tissue and cells, our study provides strong evidence to support the critical role of *AKR1B10* in smoking-associated lung cancer.

## Acknowledgments

We thank Universidade Salvador, Fundação José Silveira and Fundação Oswaldo Cruz for the support. Mr. Olival Rocha, Mr. Jose Lima, Mr. Luiz Matos, Mr. Getúlio Pacheco, Mr. Humberto and Mr. Edvan Santana (Universidade Salvador) for the technical support. This study was supported by the Intramural Program of Fundação Oswaldo Cruz (FIOCRUZ), Fundação José Silveira and by the Brazilian National Council for Scientific and Technological Development (CNPq). K.F.F. received a fellowship from the Programa Nacional de Pós-Doutorado, Coordenação de Aperfeiçoamento de Pessoal de Nível Superior (CAPES) (Finance Code 001). The work of B.B.A. was supported by grants from the NIH (U01AI115940, R01AI069923-08, R01AI20790-02). BBA and A.T.L.Q are senior investigators from CNPq. J. M. C.-A. was supported by the Organization of American States - Partnerships Program for Education and Training (OAS-PAEC) and his study was financed in part by the Coordenação de Aperfeiçoamento de Pessoal de Nível Superior - Brasil (CAPES) - Finance Code 001. M.B.A. received PhD fellowship from Fundação de Amparo à Pesquisa da Bahia (FAPESB) and FIOCRUZ. L.A.B.C. was supported by a research fellowship from CNPq. The funders had no role in study design, data collection and analysis, decision to publish, or preparation of the manuscript.

